# SubCortexMesh: A Python toolbox for surface-based analysis of subcortical brain regions

**DOI:** 10.64898/2026.09.20.753034

**Authors:** Charly H. A. Billaud, Nicolas P. M. Lavarde, Junhong Yu

## Abstract

Neuroimaging research focusing on subcortical brain regions showed that they play a role in a wide range of cognitive functions and associated their structural alterations to various psychiatric disorders. Methods to analyse such structures were developed to observe changes, not simply in terms of volumes, but in terms of surfaces, which are able to detect subtle and local changes in shape and geometrical configuration. Such advanced analyses have however been largely restricted to command line environments with relatively limited access to Python users or researchers with low technical expertise. We present SubCortexMesh as a user-friendly toolbox which covers automated surface estimation from popular subcortical volume segmentations (FreeSurfer and the *Functional Magnetic Resonance Imaging of the Brain Software Library* (FSL)), computes shape-related vertex-wise metrics (thickness, surface area and curvature) for whole cohorts and includes statistical analyses. This is all done inside Python, automatically computing all subjects of a given preprocessed directory, with the explicit intent to minimise steps and manual coding required from the users. SubCortexMesh includes statistical tools which allow conventional random field theory-based cluster analyses on native subject metrics, standardised within common surface templates, thus containing a whole workflow necessary to run an up-to-date shape-wise subcortical analyses in one package.

## Introduction

Subcortical regions of the brain play a role in a wide range of cognitive functions, including executive functions, attention, learning, language and decision-making.^1^ Structural subcortical abnormalities are prevalent in multiple psychiatric disorders including major depressive disorder (MDD)^2–4^, bipolar disorder (BD)^3,4^, schizophrenia (SZ)^3–5^, and substance dependence disorders^6,7^; and are considered to potentially play a role in pathological symptoms due to the subcortex’s role in important brain circuitry producing neurotransmitters targeted by medication^3,4^ Example of abnormalities include decreased volumes in the hippocampus, the thalamus or basal ganglia^3,4^, and have been of interest for researchers seeking to potentially improve diagnostic tools and guide treatment selection for clinicians.^3^

To study such structural changes, surface-based analyses of brain subcortical structures have emerged as an improvement over the classic volume-based approaches, which involved getting broad regional volumes (from the voxels of a MRI scan), or relying on averaged voxel-wise grey matter density (such as voxel-based morphometry^8^), which tended to overlook more focal differences that appeared in only certain parts of those regions^5,6,9^, and overall lacked information on local geometry of regional boundaries^7,10^. Surfaces, meshes made of polygons (conventionally, triangles, joined together by vertices) instead of voxels, are able to show changes that do not pertain to gross volume but to how a brain region may be thicker, thinner, expanded, contracted, at local angles and curves across the region’s shape.^10,11^

While volumetric subcortical segmentation tools have been made available in popular tools^12,13^, surface-based postprocessing of such segmentations are relatively more advanced and difficult to access for users that are not proficient in programming or are specialised in languages such as Python. To help make such analyses accessible to a broader audience of researchers, we present SubCortexMesh (SCM), a user-friendly package working with the Python programming language, for automated conversion of subcortical volumes to surfaces and computation of shape measures. SCM can run on about any computer that runs Python without requiring dockers or virtual machines.

Figure 1 depicts the standard workflow of SCM’s surface processing. It is able to read volumetric subcortical segmentations from popular automated pipelines (FreeSurfer^12^, FSL^13^). SCM aligns subject-wise segmentations to a common MNI-space template scan, extracts individual volumes for each ROI, and converts them to 3D surface meshes, including (tuneable) dilation and smoothing. SCM estimates medial curves crossing through each ROI to align them closely to a common surface template and computes surface-based metrics including thickness (radial distance from the medial curve), surface area (a third of a vertex’s triangles’ area), and mean curvature (higher curvature as concavity, lower as convexity of the surface). To standardize all meshes across subjects, the native metrics are projected to the common surface template, so each vertex can be compared across subjects. The ROIs can finally be statistically analysed separately or merged together across the whole cohort, with SCM’s slm_analysis function wrapping BrainStat’s cluster-analyses directly in Python^14^ or via VertexWiseR in *R*^15^ (similarly including BrainStat tools, as well as expanded modelling options like threshold-free cluster enhancement).

**Figure 1.**
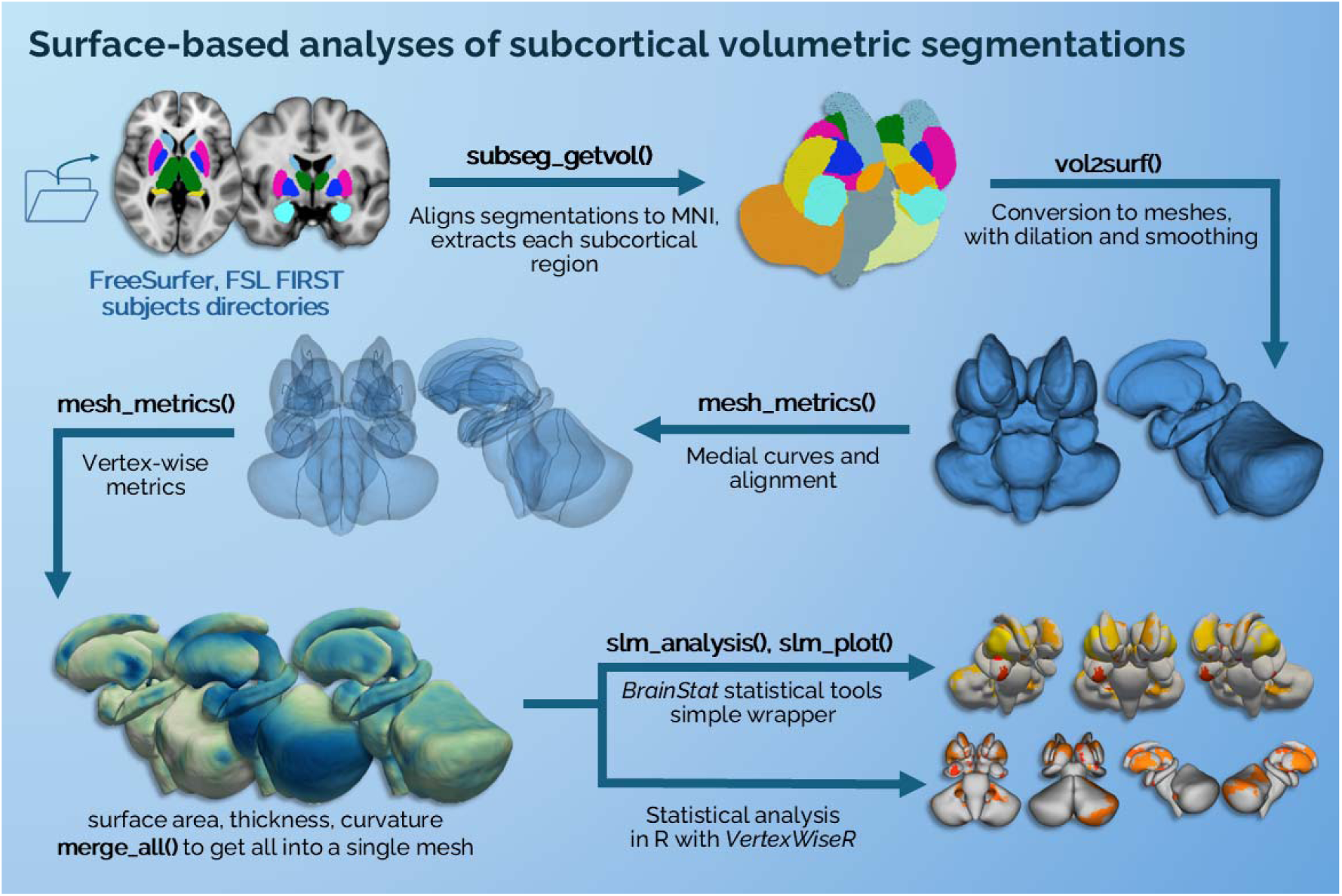
Summary of the SubCortexMesh analysis workflow.

SCM’s functionalities, along with that of software conventionally used in surface-based subcortical analyses, are presented in Table 1. SCM complements existing tools with simplified functions that minimise the amount of manual intervention and coding demanded from the users, as well as by offering fast computation (Table 3) and conventional modelling tools in Python for random field theory (RFT) cluster-based statistical analyses with any number of regions combined, in Python itself or in R via compatibility with existing statistical packages^14,15^.

**Table 1.** SubCortexMesh functionalities in parallel to existing software.

|  | SubCortexMesh | ENIGMA-shape<br>(http://enigma.usc.edu/ongoing/enigma-shape-analysis/) | FSL FIRST<sup>10</sup> |
| --- | --- | --- | --- |
| Operating system | Linux, | Linux, Mac | Linux, Mac |

|  | Windows<sup>1</sup>, Mac |  |  |
| --- | --- | --- | --- |
| **Required code environment** | Python | CLE, R, MATLAB | CLE |
| **Volume segmentations** | - | - | ? |
| **Supported segmentations** |  |  |  |
| FreeSurfer ASeg<sup>16</sup> | ? | ? | - |
| FSL FIRST (336 model)<sup>10</sup> | ? | ? | ? |
| **Additional ROIs** |  |  |  |
| Cerebellum | ? | - | ? |
| Brainstem | ? | - | ? |
| Ventral diencephalon | ? | - | - |
| **Vertex-wise measures** |  |  |  |
| Thickness (radial distance) | ? | ? | - |
| Surface area (triangles-based) | ? | - | - |
| Surface area (Jacobian-based) | - | ? | - |
| Surface area (distance-based) | - | - | ? |
| Curvature | ? | - | - |
| **Symmetrical maps** | - | ? | - |
| **Cluster-based statistics** | ? | - | ? |

Note. CLE = Command-line environment. ASeg= Automatic subcortical segmentation. ROI=Region-of-Interest. FSL FIRST= FMRIB Integrated Registration and Segmentation Tool. <sup>1</sup>Linux may also be run on Windows via a virtual machine or a subsystem, which is not required for SubCortexMesh. The volume alignment steps require a Microsoft Visual C++ redistribution library in Windows (all toolboxes include internal C++ algorithms).

As mentioned in Table 1, SCM handles FreeSurfer’s Automatic subcortical segmentation (ASeg)^16^ which is automatically outputted for each subject as part of the “recon-all” preprocessing pipeline (https://surfer.nmr.mgh.harvard.edu/fswiki/recon-all).^12^ SCM is also compatible with segmentations produced by FSL in the *FMRIB Integrated Registration and Segmentation Tool* (FSL FIRST)^13^. SCM’s reference surface templates are based on its own internal volume-to-surface conversions, applied on the fsaverage (MNI305) template’s own ASeg volume, and FSL’s training segmentation models applied to the MNI152 template^10^, including cerebellar segmentations (with 40 modes and putamen intensity as reference, as indicted by the FSL documentation).

### Package installation

SCM is compatible with Python version 3.10 and above and can be installed using Python’s

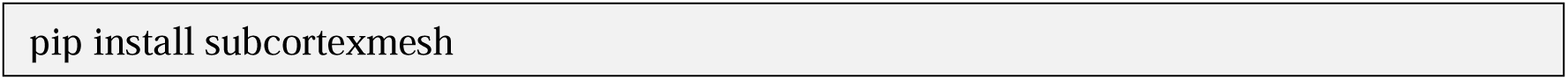

default installer: Table 2 outlines other automatically installed python tools that it depends on. In particular, SCM makes use of the *Visualization Toolkit* (VTK)^17^ an open-source 3D graphics computation software, which has been applied to clinical neuroimaging data.^18,19^ SCM uses VTK’s tools for conversion of volumes to meshes, and computation of relevant surface-based metrics, detailed later in the article.

**Table 2.** Package dependencies from other Python toolboxes.

| **SubCortexMesh function(s)** | **Related package** | **Version** | **Usage** |
| --- | --- | --- | --- |
| subseg_getvol() | AntsPy<sup>20</sup> | 0.6.2 | Segmentations-to-MNI coregistration |
| subseg_getvol(), vol2surf() | NiBabel<sup>21</sup> | 5.3.2 | .mgz and .nii volumes manipulations |
| vol2surf(), mesh_metrics(), merge_all(), autoqc_outliers() | Visualization Toolkit<sup>17</sup> | 9.5.2 | Surface conversion and manipulation, medial curves building, metrics |
| mesh_metrics(), autoqc_outliers() | Scikit-learn<sup>22</sup> | 1.7.2 | Principal component analysis to help medial curves estimation |
| mesh_metrics() | SciPy<sup>23</sup> | 1.15.2 | Rotation tools to help subject-to-template alignment |
| vol2surf(), surf_qcplot(), slm_plot(), mesh_metrics(), vis_merged(), vis_merged_flat() | Pyvista<sup>24</sup> | 0.46.4 | Interactive plots |
| surf_qcplot() | Matplotlib<sup>25</sup> | 3.10.7 | Quality check interactive plot |
| slm_analysis() | BrainStat<sup>26</sup> | 0.4.2 | Standard Linear regression models, mesh scalar smoothing |
| slm_analysis() | BrainSpace<sup>27</sup> | 0.1.22 | Mesh manipulation for use in BrainStat |

Once imported in Python, the toolbox requires base template data to be downloaded, which are the reference for standardisation of mesh metrics. Depending on the segmentations that are computed (FreeSurfer’s ASeg, FSL’s FIRST; 11.7 MB and 8.8 MB respectively), SCM will automatically prompt and assist with the template data download (in the default home path or a user-given path). No other specific operating system or environment library is required beyond the above details.

### Demonstration

For demonstration, we report an example of a study pipeline with SubCortexMesh. Validation is also provided on the same data with the ENIGMA Shape and FSL FIRST processing pipelines in Supplementary materials S1.

### Dataset

This demonstration used SUDMEX_CONN^28^, a public dataset composed of 74 cocaine use disorder (CUD) patients (31.0±7.3y, 65M) and 64 healthy controls (HC, 30.6±8.3y, 51M) (excluding 7 subjects with no group affiliation), available on *OpenNeuro* (doi:10.18112/openneuro.ds003346.v1.1.2). The data collected for the SUDMEX_CONN dataset was approved by the local ethics committee and carried out at the National Institute of Psychiatry “Ramón de la Fuente Muñiz”, Mexico City, Mexico.^28^ Previously, a study highlighted decreased surface area in the bilateral thalami and right caudate nucleus for the CUD group^7^, and we aimed to replicate similar analyses using the whole sample (the excluded 6 CUD and 12 HC in the former could not be identified upon T1 scan inspection and request).

All participants underwent a T1-weighted MRI scan using a Phillips Ingenia 3T scanner: 32-channel dS Head coil, 3D FFE SENSE sequence, TR=7 ms, TE=3.5 ms, field-of-view=240, matrix=240 × 240 mm, 180 slices, gap=0, plane=sagittal, voxel size=1 × 1 × 1 mm (five participants with voxel size=0.75 × 0.75 × 1 mm), scan time=3.19 min. The T1-weighted volumes were preprocessed using ‘recon-all’ with its default parameters in FreeSurfer 8.1.0.

### Extracting volumes and converting them to surfaces

The extraction of segmentation volumes was done on the FreeSurfer preprocessed subjects directories, automatically identifying all subjects’ segmentation volumes (aseg.mgz). For FSL, SCM would read through the run_first_all function’s output directory, looking for “*all_fast_firstseg.nii.gz” for each subject (stored in individual subdirectories).

To facilitate later standardisation, the segmentations are rigidly aligned in SCM to a common template (fsaverage/MNI305 for FreeSurfer, MNI152 for FSL) with internal use of AntsPy^20^, and then each volume saved separately per ROI. This is done for all subjects with the subseg_getvol() function (Figure 2, A). All volumes are then sequentially converted to individual surfaces using vol2surf(), which applies VTK’s marching cubes algorithm. The marching cubes approach represents the 3D voxel-wise volumes (Figure 2, B, top mesh) as triangles whose vertices “march” and intersect across the voxels to reconstruct a shape mesh (Figure 2, B, middle mesh).^29^ Here, volumes were (as per SCM’s default options) dilated and eroded to minimise graphical voxel-wise artefacts (e.g. hanging sparse voxels), and a light smoothing applied to remove artificial coarseness (such as sharp mesh spikes) of the surfaces (Figure 2, B, bottom mesh). Figure 2, C shows the interactive plots for quality control which can be launched via the surf_qcplot() function to inspect a subject’s surfaces together with their respective coregistered T1w scan. The autoqc_outliers() function can help automatically flag outliers along five global heuristic shape features (volume, surface area, sphericity, elongation, symmetry) which may then be inspected in particular with surf_qcplot(). All subject’s surfaces were visually inspected for this demonstration.

**Figure 2.**
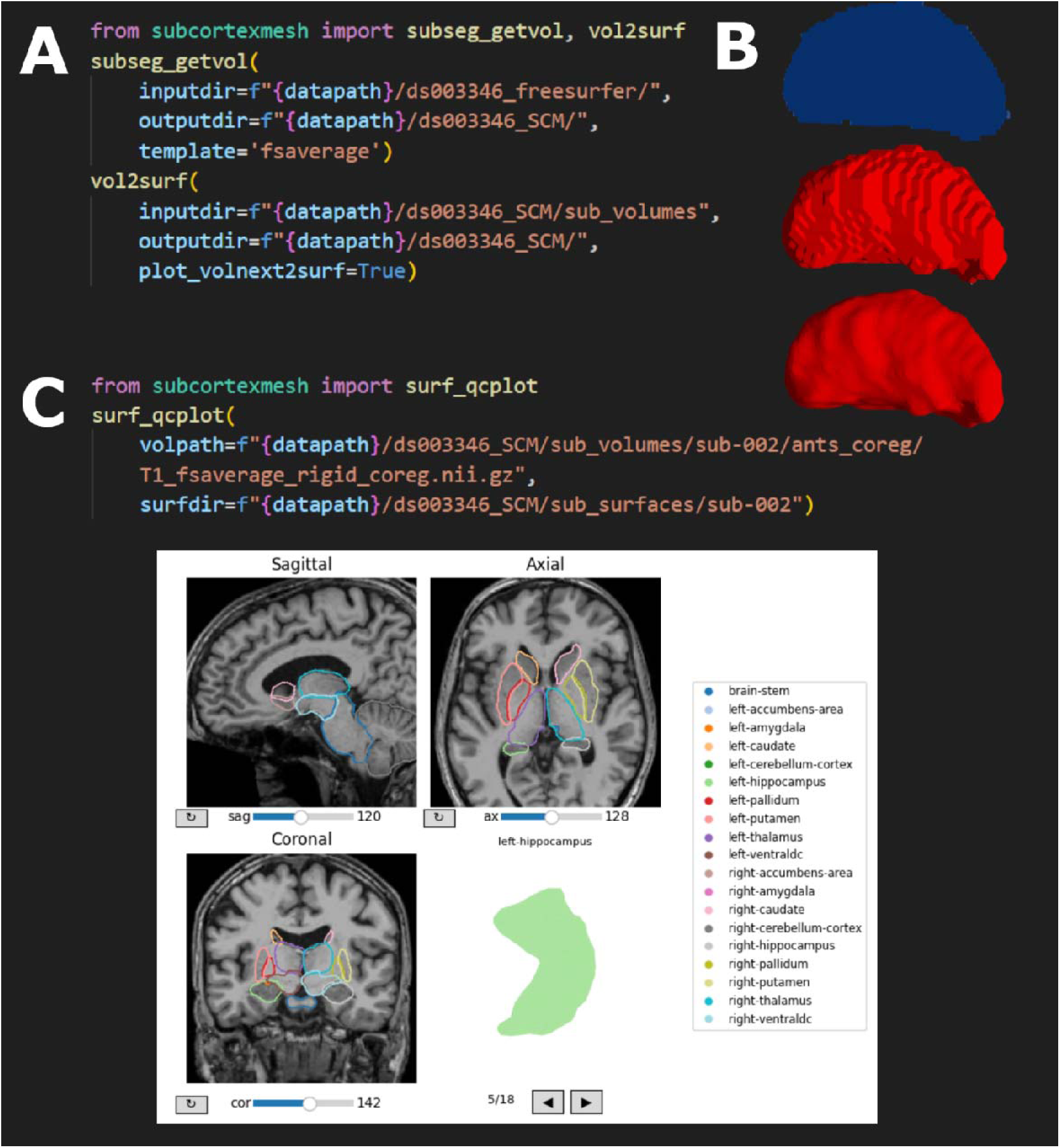
Functions to extract subcortical volumes, convert them to surfaces, and inspect them visually. *Note*. “datapath” is an example path to the directory where the FreeSurfer output data may be stored and SubCortexMesh’s output directories go. A) Functions to extract subcortical volumes of all subjects of a preprocessing output directory and to convert them to surface meshes. B) Output of subseg_getvol() and vol2surf() for a subject’s left thalamus: extracted volume (top), volume converted to a surface via marching cubes (middle), and the final smoothed mesh (bottom). vol2surf()’s *plot_volnext2surf* option plots the final mesh next to its original voxel-wise volume. C) function to visually inspect surface meshes of a subject on top of their respective scan and screenshot of its interactive window. The dilation and smoothing by default to minimise graphical artefacts will make the boundaries appear slightly wider than their original anatomy, but the plots can be viewed on surfaces produced with dilate_erode=False in vol2surf().

### Estimating surface-based metrics

In a similar fashion to the ENIGMA shape pipeline^11^, SCM estimates a medial curve inside each surface, which is a line crossing through the middle of a shape, as part of the mesh_metrics() function (Figure 3, A). The medial curve has two roles in SCM: calculating surface thickness (described further below, depicted in Figure 3, B), and optimising the alignment between the subject-space meshes and the common template meshes (which also have their own medial curves). In SCM, the medial curve is obtained by identifying the overall directional axis across the mesh (i.e., the direction across which its vertices extend the most), and “slicing” the mesh in one hundred planes from both ends of the axis. The curve is defined as a line traced across the centroids of each plane, linking both of their ends and reduced to twenty-five subdivisions across the line to “smooth” its path.

**Figure 3.**
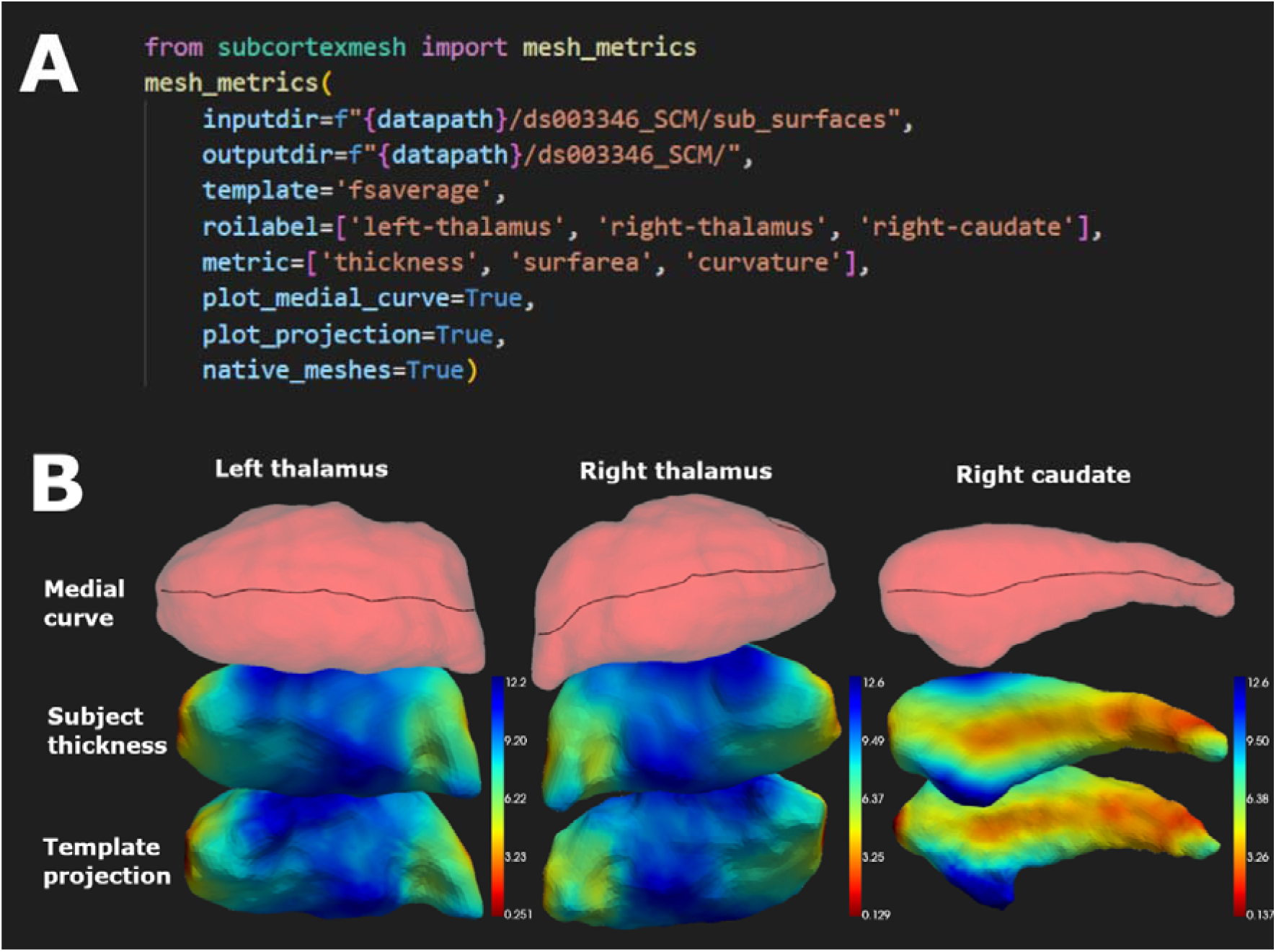
Function to compute surface-based metrics and project them to a standard space. *Note*. “datapath” is an example path to the directory where the SCM surface output directories may be stored. A) Function to compute, for all subjects, a medial curve inside each selected mesh, as well as calculating selected vertex-wise metrics and projecting them to a standard template surface. B) Each mesh shows a stage of mesh_metrics()’s processing, including medial curve estimation (top mesh), subject-wise metrics calculation (thickness displayed here in the middle mesh), and projection to a common surface template (bottom mesh).

The thickness measure is the distance separating each vertex on the surface from the closest point on the medial curve, or *radial distance*, and is computed with VTK’s *vtkImplicitPolyDataDistance* function (Figure 3, B). Additionally, two vertex-wise measures are computed by default by SCM: surface area, which follows FreeSurfer’s own measure as 1/3 of the sum of the area covered by the triangles that a vertex belongs to; and mean curvature (VTK’s *vtkCurvatures*, inverted to match FreeSurfer’s convention), which indicates how bent is the surface at a vertex, with higher curvature meaning more concave surface and lower curvature a more convex surface. We reported correlations between the metrics for the same vertices in Supplementary material S2, showing that there was little collinearity between SCM’s surface-based measures (maximum *r*=0.21).

While the mesh_metrics() function gives the option to keep native meshes (*native_meshes* argument, Figure 3, A), all native vertex-wise values are projected to a common surface template (here, based on fsaverage, Figure 3, B, middle-to-bottom mesh shows a subject-to-template projection). The “projection” consists in first aligning the subject mesh to the template through closest alignment of their medial curves, applying a transform to make the subject mesh’s vertices match as closely as possible the template’s (VTK’s iterative closest points algorithm with rotation, translation and isotropic scaling) and finally assigning values in the template to vertices that are the closest to each other (using VTK’s *vtkPointLocator*).

### Statistics

Using SCM’s slm_analysis() function for statistical modelling, a simple fixed effects linear model (wrapping BrainStat’s *SLM*^26^) was run, testing a simple binary group difference, with no covariates, between CUD and control groups (Figure 4, A), to replicate Xu and colleagues’ previous use of the vertex-wise analysis in FSL FIRST^7^. Three models were run, one for each metric (thickness, curvature, surface area) on the three selected ROIs together. RFT cluster correction was applied with a significance threshold at *p* < .05 (Figure 4, B).

**Figure 4.**
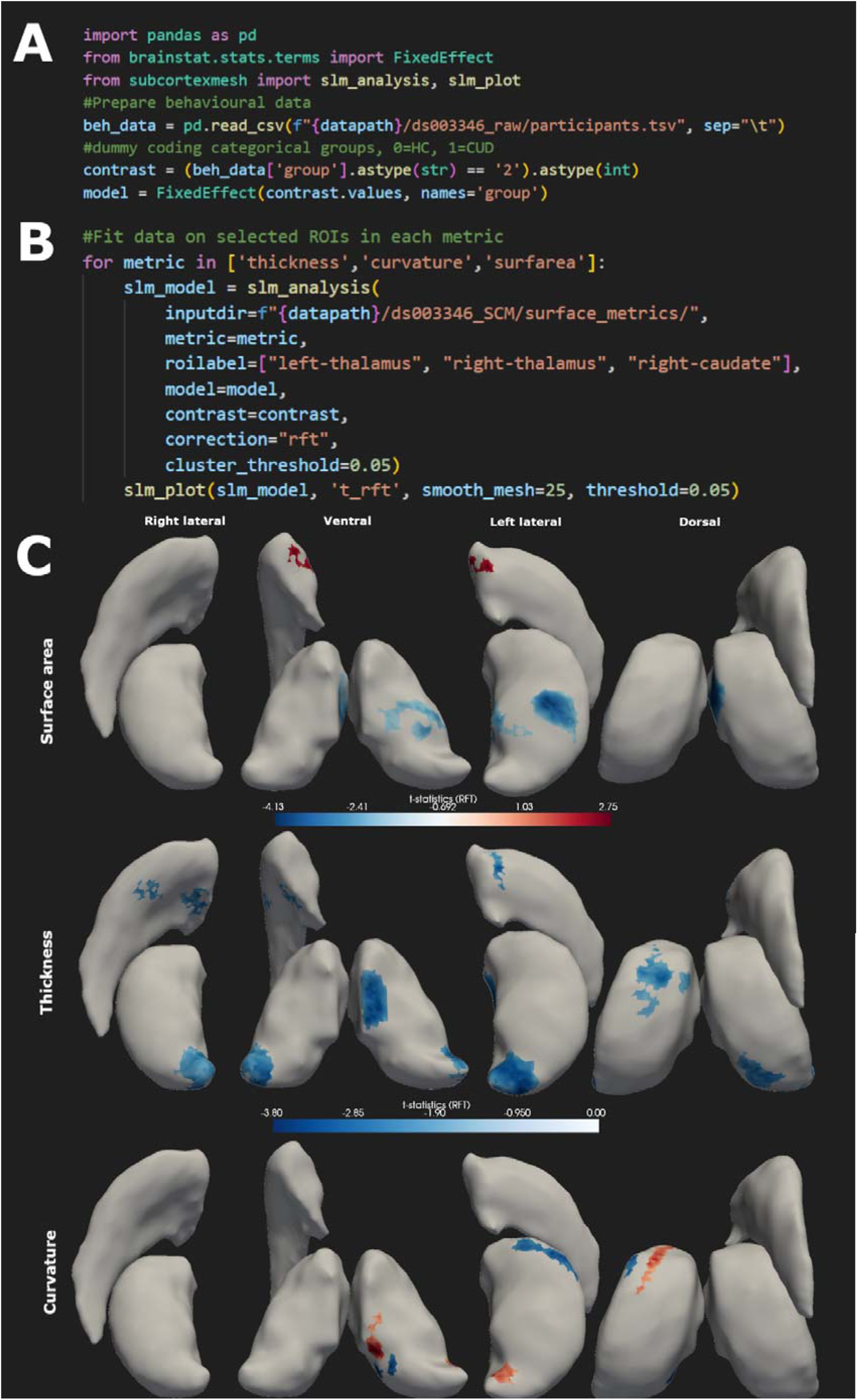
Functions to compute statistical analyses with the surface-based metrics. *Note*. RFT= Random field theory. A) Example of behavioural data, with ‘contrast’ containing the main effect of interest (group, dummy coded) and ‘model’ the contrast along with its (potential) control variables, formatted for BrainStat’s fixed effects models. B) SubCortexMesh’s function slm_analysis() to test the effect of the contrast variable on the vertex-wise values of selected regions of interest, with RFT cluster-correction (other option is ‘fdr’). slm_plot() is a function which can plot different maps from the fitted model: here, the *t*-statistics map filtered with cluster-wise significance. C) the vertex-wise t-values overlayed on the bilateral thalamus and right caudate, thresholded form the significance of clusters they belonged to (at *p*<.05), for each metric.

SCM also automatically outputs tables of descriptive statistics, including average vertex-wise values, standard deviations, minimum, maximum and range of values, for each ROI of a subject in native-space. Additionally, to investigate whether these ROI-level metrics were significantly different across groups, independent *t*-tests across groups were run for the same ROIs and metrics, with correction for false discovery rate (FDR).

## Results

### Vertex-wise effect of group on surface-based metrics

Figure 4, B shows the significant clusters where there was an effect of group in each metric after applying RFT cluster-correction. In the CUD group compared to the control group, the left thalamus had lower middle ventral and lateral surface area, lower thickness in ventral/medial and posterior lateral parts, lower curvature in the anterior lateral and posterior ventral parts, and higher curvature in anterior dorsal and posterior medial/ventral parts. The right thalamus had lower anterior medial surface area, and lower posterior medial and lateral thickness. The right caudate had lower thickness in the anterior medial and lateral parts, and higher surface area in the anterior ventral part.

Decreases in the same ROIs could be found using ENIGMA Shape and FSL FIRST (Supplementary material S1), although no significant vertices survived ENIGMA’s regional searchlight FDR correction, while no cluster reached significance for FSL FIRST’s right caudate (minimum cluster-corrected *p*-value was 0.08). There was mixed spatial overlap between the significant clusters, and patterns of decreases most resembling SCM’s were found in the left thalamus (posterior tip, medial side) in both the ENIGMA and FSL FIRST outputs, FSL FIRST’s output for the right thalamus (posterior parts), and ENIGMA’s right caudate (anterior parts) outputs. These difference in results may be explained by the difference in the templates’ shapes themselves, but also the different metric computation approaches (especially the measures of surface area and the curvature measure which all differ from each other conceptually).

### Group differences in average surface-based metrics

Figure 5 reports the group differences in average ROI-wise values for each metric computed by SCM. Only the left and right thalami’s surface area were significantly lower on average in the CUD group after FDR correction.

**Figure 5.**
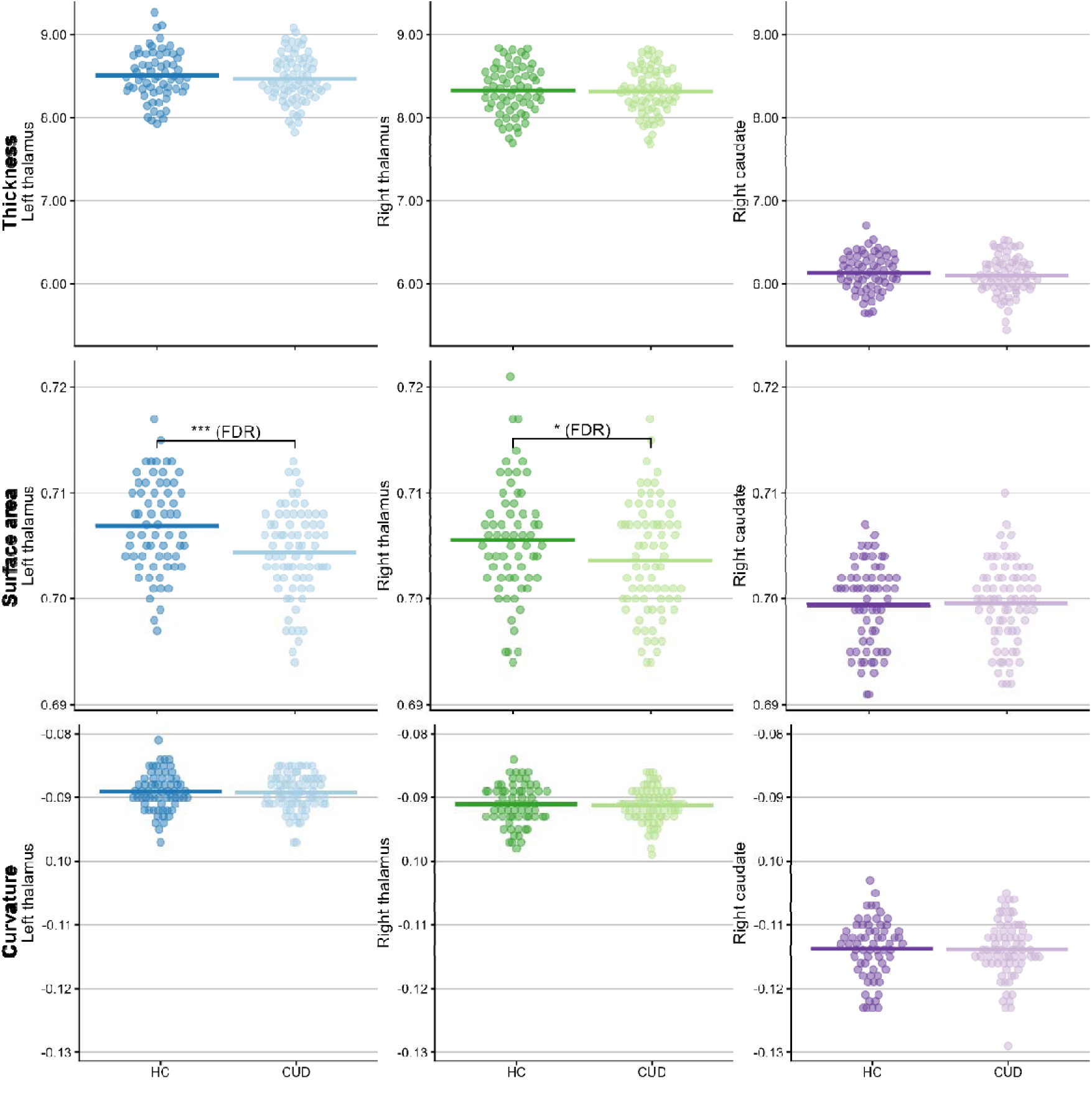
Group differences in average vertex-wise metric per region-of-interest. *Note*. HC=Healthy controls. CUD= Cocaine user disorder. FDR= (survives adjustment for) false discovery rate. * Significant at *p*<.05. *** significant at *p*<.001.

Significant group differences in ROI-wise average metrics could not be found with ENIGMA Shape or FSL FIRT after FDR correction, but the thickness averages of SCM and ENIGMA’s correlated strongly (further details and discussion in Supplementary material S1).

### Analysing all surfaces at once

SCM allows all surfaces to be merged and analysed at once with the merge_all() function, which gather metrics data into one global surface object whose vertices are all tested and corrected together (Figure 6, A). For this type of data slm_plot() gives the option to plot the ROIs in an interactive 3D viewer, with a slider to space out the regions and inspect them at 360° (Figure 6, B), or in a 2D grid-like image with pairs of regions in top and bottom views (Figure 6, C). Applying this to the same demo model, as an exploratory example, highlights additional group differences in the thickness of the right hippocampus, right putamen, bilateral cerebella and ventral diencephala.

**Figure 6.**
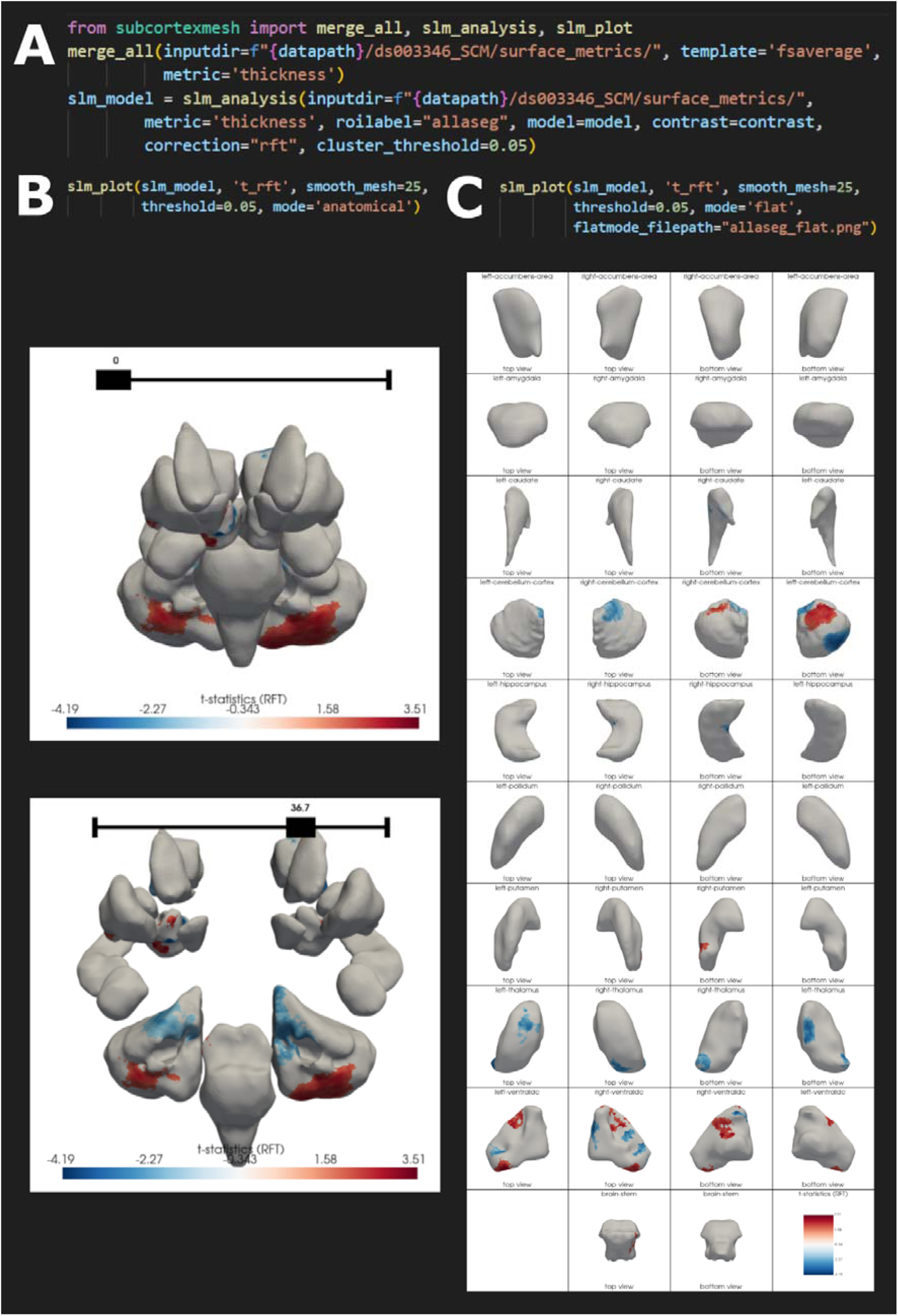
Merging all subcortical regions for analysis and plotting. *Note*. RFT= Random field theory. A) Function merging all surface outputs for a given metric produced by mesh_metrics(), and the same model being run on the resulting “allaseg” global surface object (named “allfslfirst” for the FSL FIRST template). B) 3D interactive plotter which allows 360° inspection, with a slider that determines how spaced out are the regions from each other, to facilitate visualisation. C) 2D plotter which prints a grid where all regions are laid out separately, for both hemispheres, with top views and bottom views.

### Completion times

Table 3 details average computation times, per subject, assuming all surfaces are to be computed from a FreeSurfer preprocessing output, including all metrics, with two different systems. The coregistration done with ANTsPy uses multithreading by default, and specific numbers of threads can be set by modifying the environment variable before importing ANTS in Python (e.g.:*os.environ[“ITK_GLOBAL_DEFAULT_NUMBER_OF_THREADS”]=“8”).* VTK kept the same sequential, 1 thread usage and its processing speed may vary with the performance of the CPU system. Note that while the functions run subjects sequentially, they can still be run in multiple parallel processes without conflict, as each subject will have a 1-hour-old temporary file marking it as already running.

**Table 3.**
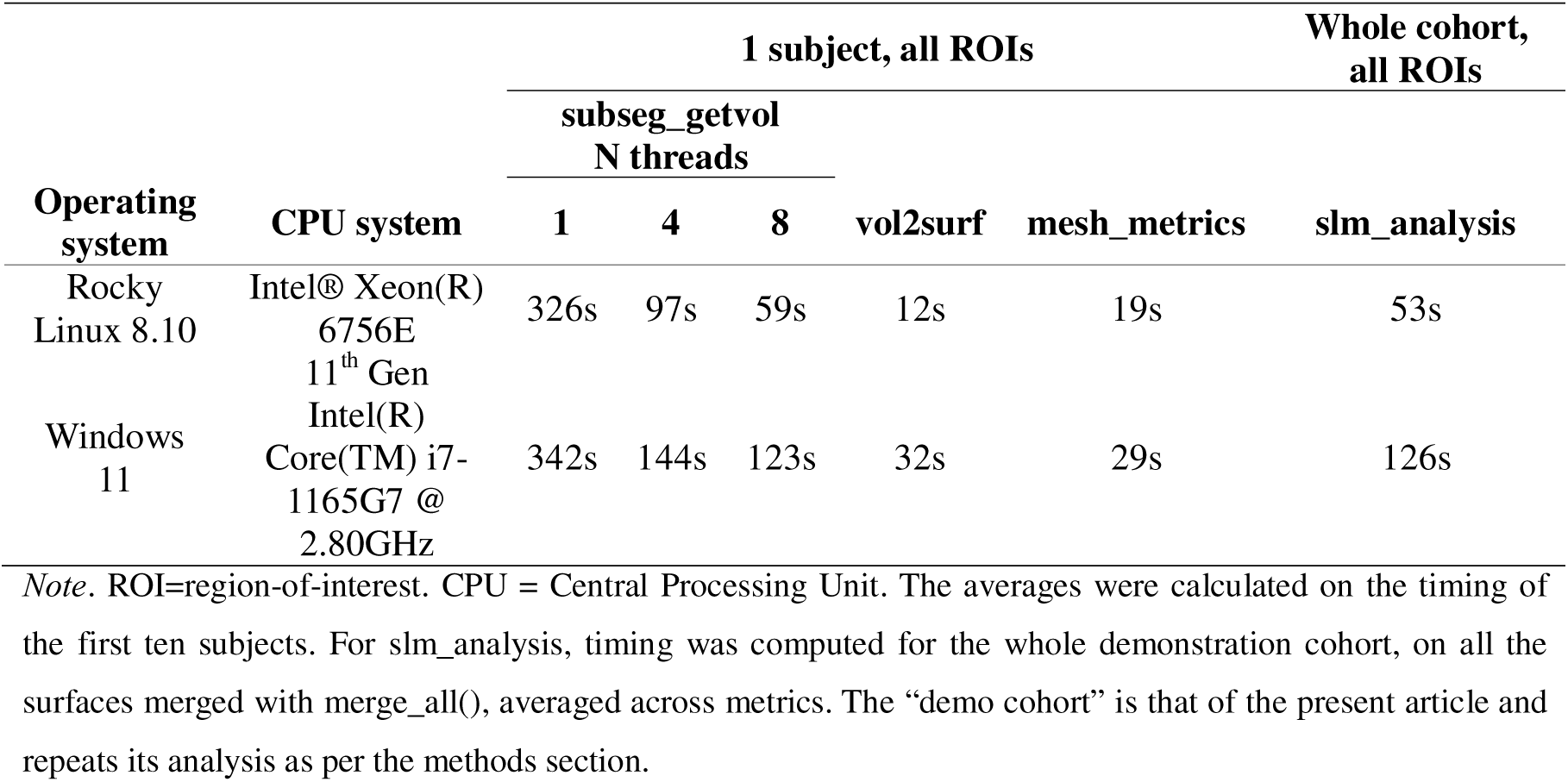
Average processing durations across SubCortexMesh’s pipeline stages, depending on the number of allocated threads.

## Discussion

SubCortexMesh is a toolbox that helps researchers implement surface-based analyses of subcortical brain regions in Python, and takes advantage of available graphical tools (Table 2) to apply them on outputs from popular volumetric segmentations algorithms (FreeSurfer^16^ and FSL^10^). SCM offers a streamlined pipeline (Figure 1) that guides users from the volume-to-surface conversions, through surface-wise metrics computation, up to conventional statistical modelling application, in an automated fashion with minimal system requirements and manual coding.

SCM reproduces a radial-distance based measure of thickness, inspired by the ENIGMA Shape approach^11^, and complements other tools with the inclusion of curvature and triangle-based surface area measures (which follows FreeSurfer’s own concepts). It also applies them to non-standard surface-based structures such as the cerebellum, brain-stem and the ventral diencephalon from the ASeg parcellation scheme^16^. The processed data from SCM’s surface-based pipeline can then be analysed with statistical tools in R^15^ or Python to flexibly apply analyses on the surface vertices (one at a time, or grouped together with any number of available subcortices).

As a demonstration, we outlined the steps of the SCM toolbox by replicating a previous vertex-wise analysis run on a public dataset using FSL FIRST^7^. As in the previous study, SCM did report significant shape alterations, including thinning and reduced surface area, in the regions-of-interests that the previous study had identified with decreased surface area in a group with cocaine use disorder. This reinforced the hypothesis that surface-based alterations of the thalamus and caudate nucleus were tied to substance addiction, given their roles in the dopaminergic pathways and addictive behaviours.^7^ SCM also added varying alteration patterns in the thalami’s shape with the introduction of the curvature metric, highlighting a form of potential neural reshaping, further highlighting subtle geometric changes that voxel-wise volumetric analysis would normally miss.

SCM fills a gap in the availability of tools dedicated to such approach in Python, and opens the door to researchers who are more familiar with the Python language to also run shape analysis of the subcortex, and makes it especially simple for beginners who might not be comfortable manually modifying code, file sheets and/or switching environments. There are still limitations that remain to be addressed and could potentially be part of future updates, such as its limitation to only two subcortical segmentation schemes, its current lack of automated quality assessment, or the unavailability of subfield or subnuclear mesh parcellations which could help researchers interpret the location of their observations from an anatomical standpoint.

## Data and Code Availability

The SUDMEX-CONN neuroimaging dataset is available on OpenNeuro: https://doi.org/10.18112/openneuro.ds003346.v1.1.2

The code produced to develop SubCortexMesh is made publicly available on its git repository at https://github.com/chabld/SubCortexMesh

## Declaration of generative AI use

Anthropic’s *Claude* (claude.ai) large language model (LMM) was used during the development of the software’s code, for assistance such as function optimisation and error proofing. No package function relied blindly on a prompt, all lines of code generated by the LLM, as well as their outputs, were manually inspected, verified and adapted by human examiners. The article itself was not written using any LLM.

## CRediT authorship contribution statement

**CHAB**: Conceptualization; Data curation; Formal analysis; Software; Validation; Investigation; Visualisation; Methodology; Writing – original draft; Writing – review & editing. **NPML**: Software; Visualisation. **JY**: Supervision; Funding acquisition; Project administration; Resources; Writing – review & editing.

