## Supplementary materials for "SubCortexMesh: A Python toolbox for surface-based analysis of subcortical brain regions"

### S1. Validation of the subcortical analysis pipeline using other toolboxes

In the following sections, we repeated the same analyses from the main text using the post-processed data from the the FSL FIRST^1^ pipeline and ENIGMA Shape^2^ pipeline (http://enigma.usc.edu/ongoing/
enigma-shape-analysis/).

### Demonstration

#### Surface processing

ENIGMA Shape was used on the same FreeSurfer preprocessing output as SCM. Meshes were drawn from the segmentations with topological correction and smoothing, registered to a standardised surface template via “medial demons” registration, involving spherical inflation and matching local geometric features^2,3^. ENIGMA Shape’s template is derived from an Euclidean average of 200 (100 males) participants from the Queensland Twin Imaging Study (QTIM) ^3^. Two measures were calculated: thickness, as radial distance from estimated medial curves, and surface area as the log of the Jacobian determinant, which represents the ratio of the subject-wise shape vertices relative to the matching vertices in the template-wise shape^3^. Each unilateral region’ measures were kept separately (instead of averaged or mirrored across hemispheres).

FSL FIRST applies its own segmentations on individual T1w volumes, with subject meshes rigidly aligned and scaled to the FSL training model’s mean shape^1^. The training model is derived from a dataset of 336 manually labelled T1w images composed of normal and pathological brains^1^. For each subject mesh, the “surface area” is measured as the vertex-wise perpendicular distance relative to the cohort-wise average shape. A negative value means the subject’s vertex moves inwardly, interpreted as atrophy, while a positive value means the vertex moves outwardly, interpreted as expansion^1^.

#### Statistics

Since toolboxes differed in their statistical modelling approaches, each toolbox’s respective standard methods were used, as closely as possible to previous approaches on the dataset^4^. All of the models tested a simple binary group difference, with no covariates, between CUD and control groups on each ROI separately: left thalamus, right thalamus, and right caudate.

For the ENIGMA Shape pipeline, linear regression models at each vertex (*mass_uv_regr_*_uni.sh*) were used, testing for the effect of group. Outputted p-value maps were corrected using searchlight false discovery rate^5^, a Gaussian weighting function with 6 mm full-width half maximum (FWHM) kernel size for the searchlight and a 2 mm Gaussian smoothing^3^.

For the FSL FIRST modelling, a general linear model was used (*randomise* function) with the default 5000 nonparametric permutations of residuals. The t-statistics contrasts were corrected for multiple comparison with cluster-based family-wise error rate (FWE) at *p* <.05 using the null distribution of the maximum cluster mass across the image (-C flag, cluster-forming threshold at 1.96).

### Results

##### Vertex-wise effect of group on surface-based metrics

Figure S1 shows the vertex-wise findings for each toolbox, ROI, and available metric. To illustrate the data trends across the meshes, VertexWiseR^6^’s transparent thresholding method for overlaying plots was used to report the significant clusters of SCM as well as non-significant t-values. None of the ENIGMA pipeline results survived correction via Langers searchlight FDR correction (using Gaussian weighting function with 6 mm full-width half maximum (FWHM) kernel size and 2 mm Gaussian smoothing following^3^), and the plot displaying beta values at *p<*.05 (uncorrected). In the FSL FIRST pipeline, only the left and right thalami had significant clusters of decreases in CUD. This latter difference might be explained by the previous study’s excluded participants^4^ whom we could not identify (we tried reaching out to the previous authors to know the IDs of those who were excluded exactly, but did not receive an answer).

**Figure S1. Group differences in surface-based metrics across toolboxes**

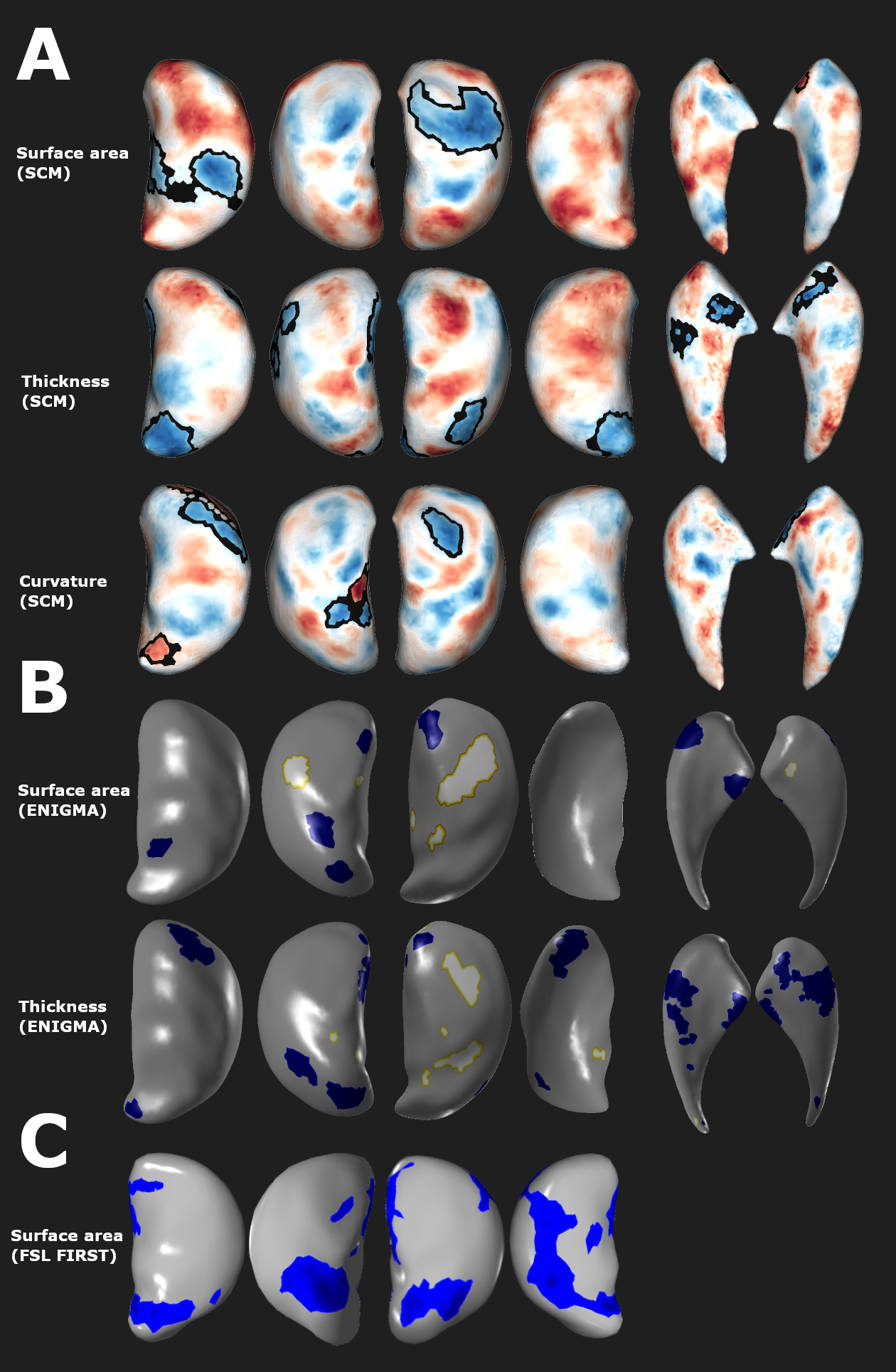
*Note*. A) Thalamus surface plots showing the t-statistics for the group effect on surface area, thickness and curvature, with significant vertices with random field theory cluster correction (at *p*<.05) outlined with a black boundary (overlaid on non-significant t-statistics using the VertexWiseR package in R). B) Vertex-wise results with the ENIGMA Shape pipeline. Thalamus surface plotsshowing the beta values for the group effect on Jacobian surface area and thickness, significant at *p* < .05 (none survived searchlight FDR correction). For the sake of visualisation, ENIGMA surfaces have been flipped as the meshes are mirrored in their individual templates (e.g. right caudate pointing the opposite way). C) Vertex-wise results with the FSL FIRST pipeline. Thalamus surface plots showing the t-statistics for the group effect on distance-based surface area, with permutation-based cluster mass-correction at *p*<.05. The right caudate data is missing for FSL FIRST as no significant (minimum cluster corrected *p*-value was .08).

##### Group differences in average surface-based metrics

Group differences in metrics across pipelines are depicted in Figure S2. The average surface-based metrics were computed for each region-of-interest in all subjects with SCM as per the main article. This was repeated with the ENIGMA Shape output (by averaging the vertex-wise values in each subject’s output). For FSL FIRST, the vertex-wise results of subjects are projected to a single 4D volume space, and the non-empty voxel-wise values corresponding to the vertex-wise values are averaged per subject (masking out the background voxels).

Results show that most ROI-wise average metrics could not detect a significant group difference, except for SCM’s average surface area, significantly decreased in the CUD group; and FSL’s distance-based surface area (relative to the average cohort-based mesh), which was increased in the CUD group in the right thalamus and the right caudate. FSL’s counter-intuitive finding is arguably the result of the dilution of the signed values in the averaging, with overall weaker or less consistent increases in the ROIs’ shape surface in CUD, despite stronger and more consistent clusters that were significantly lower in the CUD group, which shows ROI-wise values to be harder to interpret than vertex-wise specific patterns. Only the SCM findings survived FDR correction.

**Figure S2. Group differences in average vertex-wise metric per region-of-interest and toolboxes**

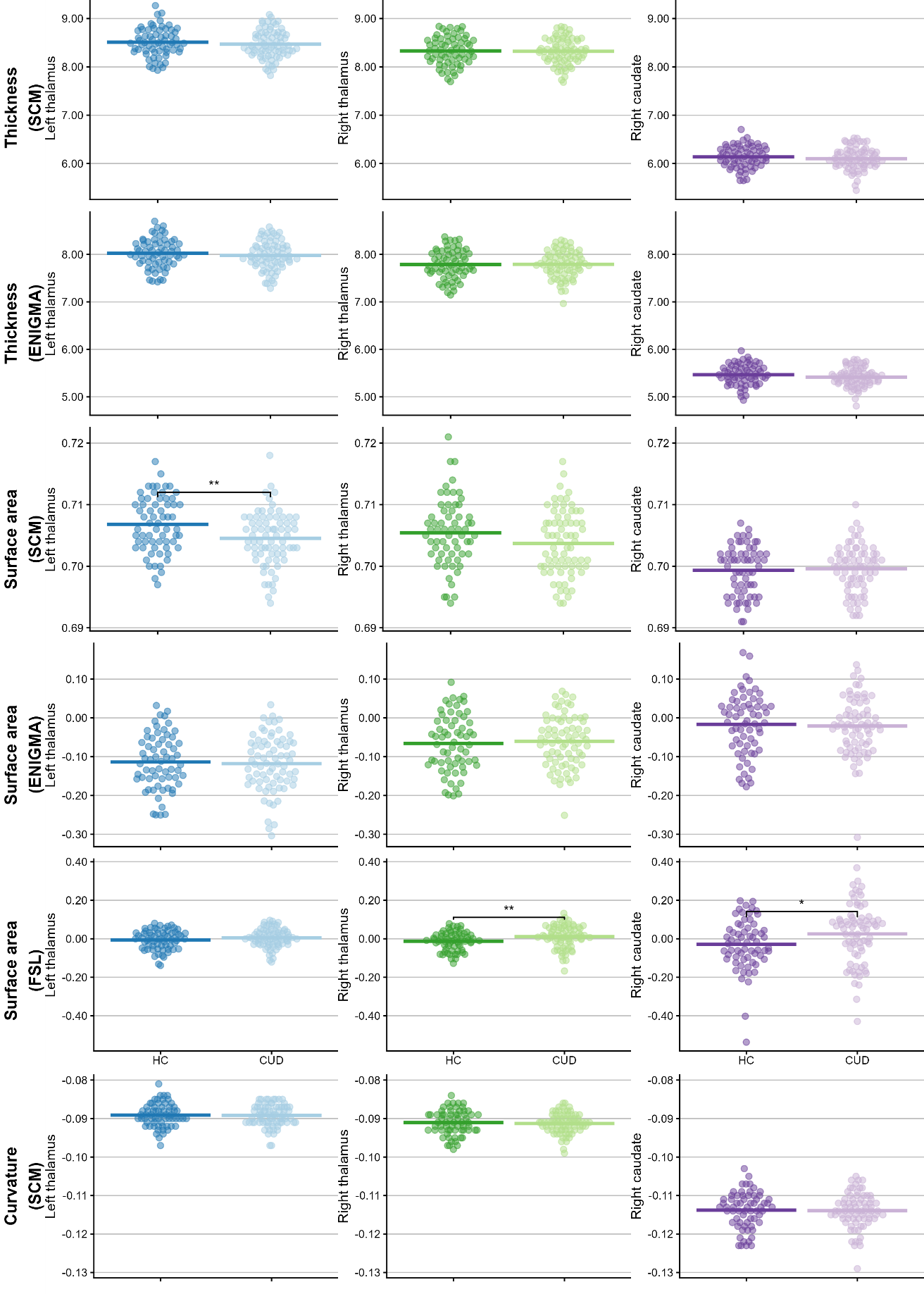

*Note*. HC=Healthy controls. CUD= Cocaine user disorder. FDR= (survives adjustment for) false discovery rate. SCM=SubCortexMesh. FSL= FMRIB (Functional Magnetic Resonance Imaging of the Brain) Software Library. * significant at *p*<.05. ** significant at *p*<.05. *** significant at *p*<.001.

Correlations across the toolboxes’ average metrics are illustrated in Figure S3. Both SCM and ENIGMA’s respective thickness measures correlate strongly (0.9), which confirms their broad similarity. The slightly thicker SCM averages, as visible in Figure S2, may be explained by the dilation of the surfaces that SCM operates by default to correct for artefacts, while the ENIGMA shape pipeline relies on spherical deformation which may not require it^2^. SCM’s triangle-based surface area correlates negatively and/or very weakly to the rest of the metrics, indicating that triangle-based surface area is specifically distinct from other measures, although it is most sensitive to group differences and thus a relevant feature (Figure S2). FSL’s distance-based surface area similarly only weakly correlated with other measures.

**Figure S3. Correlations across average surface-based metrics and toolboxes**
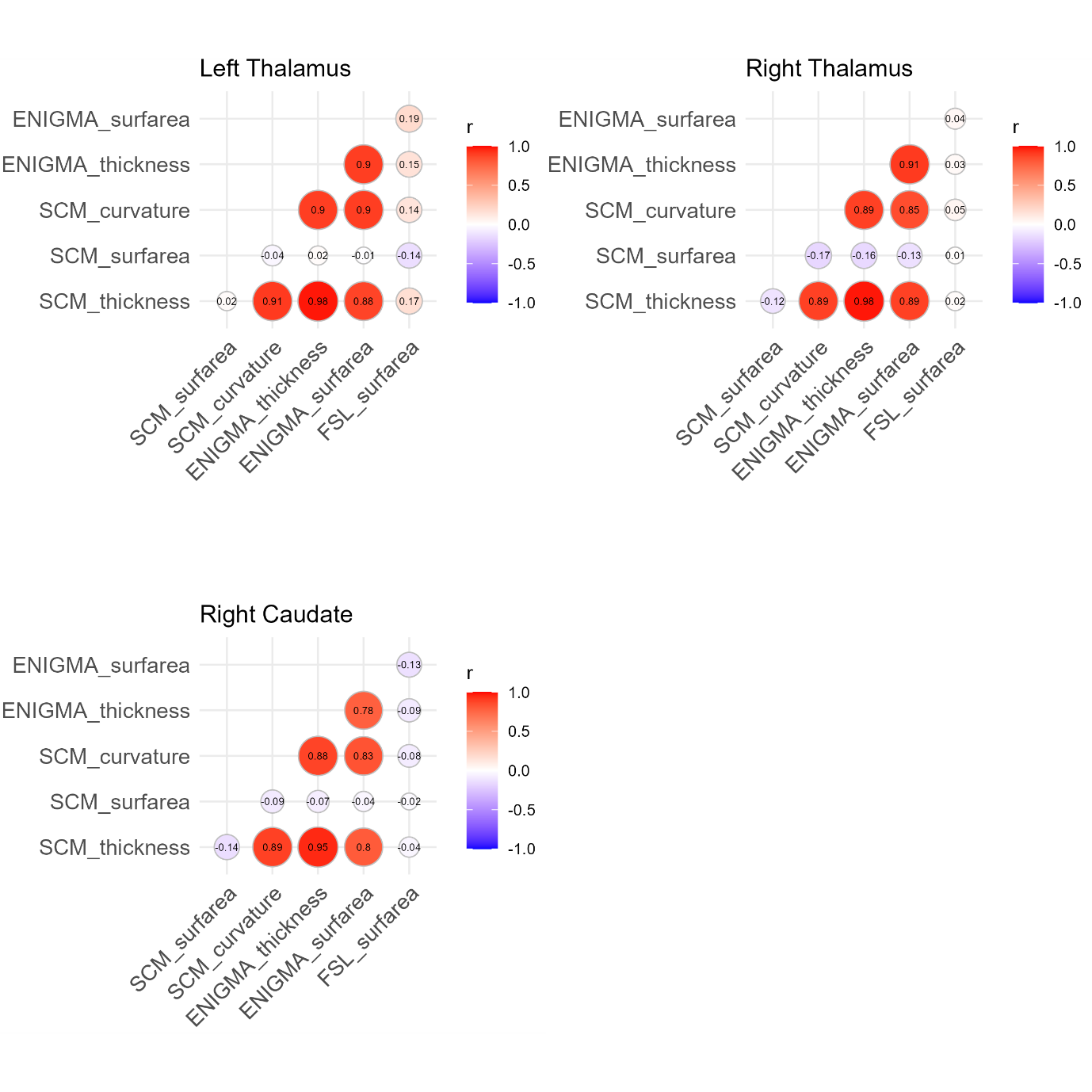

*Note*. SCM=SubCortexMesh. FSL= FMRIB (Functional Magnetic Resonance Imaging of the Brain) Software Library.

### S2. Vertex-wise correlations between SubCortexMesh’s surface-based metrics

Figure S4 reports Pearson correlations between the average metric values of each vertex (averaged across the cohort) and the average values of the other metrics for the same vertices. It shows that SCM’s measures are very distinct, with very low to low *r* values and the strongest correlations being between curvature and thickness.

**Figure S4. Correlations between vertex-wise average metrics across the whole cohort**
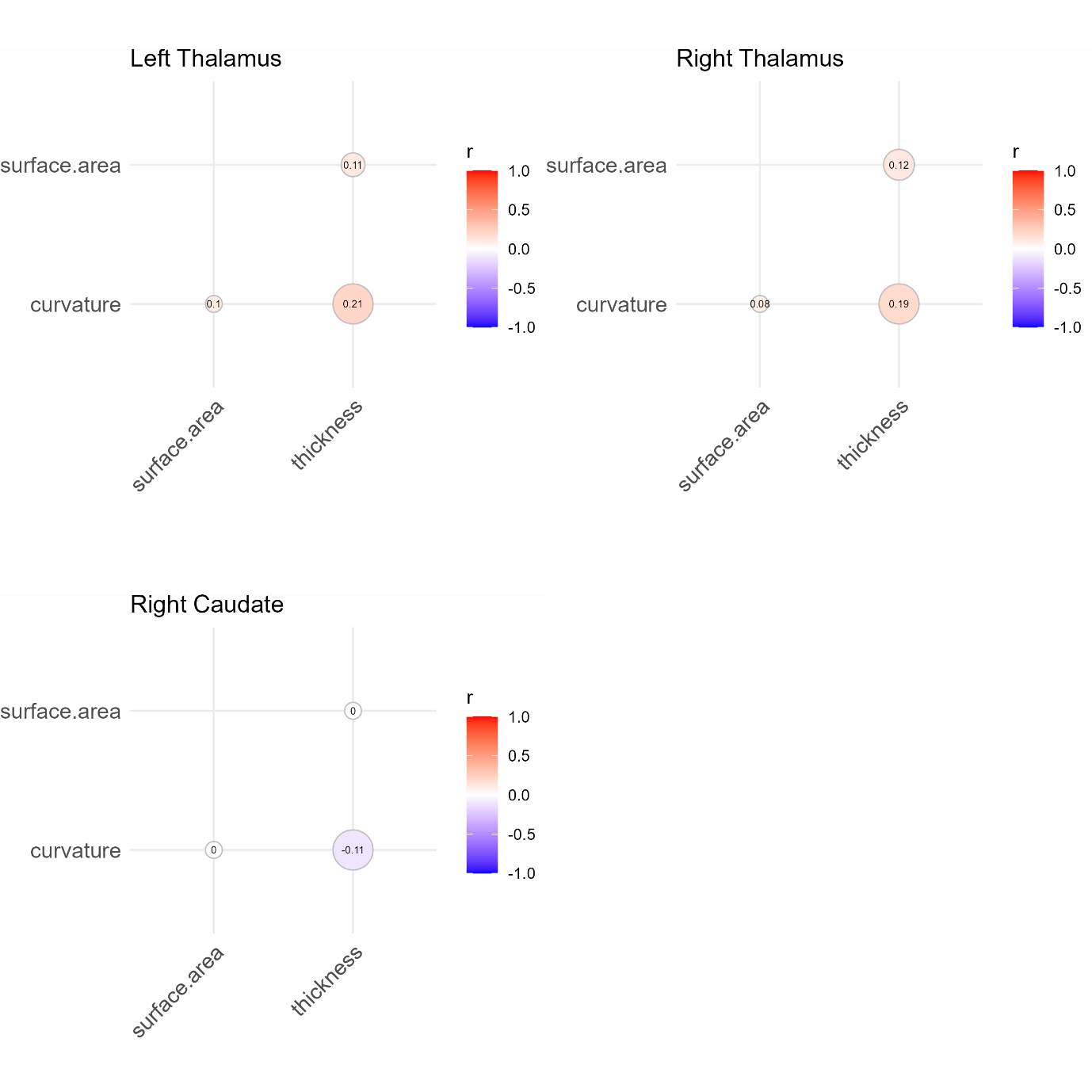
